# Reciprocal polarization imaging of complex media

**DOI:** 10.1101/2023.05.19.541234

**Authors:** Zhineng Xie, Guowu Huang, Weihao Lin, Xin Jin, Xiafei Qian, Min Xu

**Author notes:** Correspondence: Min Xu.

## Abstract

The vectorial evolution of polarized light interaction with a medium can reveal its microstructure and anisotropy beyond what can be obtained from scalar light interaction. Anisotropic properties (diattenuation, retardance, and depolarization) of a complex medium can be quantified by polarization imaging by measuring the Mueller matrix. However, polarization imaging in the reflection geometry, ubiquitous and often preferred in diverse applications, has suffered a poor recovery of the medium’s anisotropic properties due to the lack of suitable decomposition of the Mueller matrices measured inside a backward geometry. Here, we present reciprocal polarization imaging of complex media after introducing reciprocal polar decomposition for backscattering Mueller matrices. Based on the reciprocity of the optical wave in its forward and backward scattering paths, the anisotropic diattenuation, retardance, and depolarization of a complex medium are determined by measuring the backscattering Mueller matrix. We demonstrate reciprocal polarization imaging in various applications for quantifying complex non-chiral and chiral media, uncovering their anisotropic microstructures with remarkable clarity and accuracy. Reciprocal polarization imaging will be instrumental in imaging complex media from remote sensing to biomedicine and will open up new applications of polarization optics in reflection geometry.

## 1 Introduction

The interaction of light with a medium provides a non-invasive means for characterization and imaging. In addition to intensity, phase, coherence, and spectrum variations of scalar light[1], how the vector wave evolves when interacting with a medium for polarized light can reveal the microscopic structure and anisotropy of the medium[2]. The use of polarization optics has recently been expanding rapidly in biomedicine[2-6] and applied in, for example, the characterization of complex random media[7,8], tissue diagnosis[9,10], advanced fluorescence microscopy[11], and clinical and preclinical applications[4,12]. The polarized light interaction with media is succinctly described by the four-by-four Mueller matrix ***M***, which transforms the state of polarization (the Stokes vector) of the incident beam to that of the outgoing beam[13]. The medium microscopic structure is encoded in the sixteen elements of the Mueller matrix without a direct link. It requires, therefore, decomposition to interpret the polarized light-medium interaction and link the evolution of the vector light wave to the physical property of a complex medium. The standard polar decomposition (or the Lu-Chipman decomposition[14]) factors the Mueller matrix ***M*** into a product of a depolarizer matrix ***M***_Δ_, a retarder matrix ***M***_*R*_, and a diattenuator matrix ***M***_*D*_ where the individual matrices describe anisotropic depolarization, phase, and amplitude modulation of polarized light by the medium, providing a straightforward phenomenological interpretation of basic optical properties (depolarization, birefringence, and dichroism) as well as the underlying microstructure of the medium. Variants of the polar decomposition[15,16] and differential Mueller matrix decomposition[17,18] have also emerged recently. However, no existing decomposition method suits the analysis of the Mueller matrices measured inside a backward geometry, which is ubiquitous and often preferred in diverse applications from remote sensing to tissue characterization. In reflection geometry, the probing polarized light traverses the sample along the forward and backward paths in sequence. More importantly, the anisotropic polarization property of the sample seen by the probing beam in the forward and backward paths is reciprocal to each other and not the same in general.

In this article, we present reciprocal polarization imaging of complex media by introducing reciprocal polar decomposition of backscattering Mueller matrices (see Fig.1). In contrast to the Lu-Chipman decomposition of the Mueller matrix into the product of ***M***_Δ_ ***M***_*R*_ ***M***_*D*_, the reciprocal polar decomposition factors the backscattering Mueller matrix into a product of 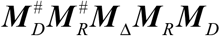 with the diattenuation and retardance along the backward path specified by the reciprocal of their counterparts 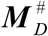 and 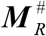 along the forward path based on the reciprocity of the optical wave in its forward and backward paths. We then demonstrate reciprocal polarization imaging in various applications for quantifying complex non-chiral and chiral media, uncovering their anisotropic microstructures with remarkable clarity. Reciprocal polarization imaging will be instrumental in imaging complex media and opening up new applications of polarization optics in reflection geometry.

**Fig. 1.**
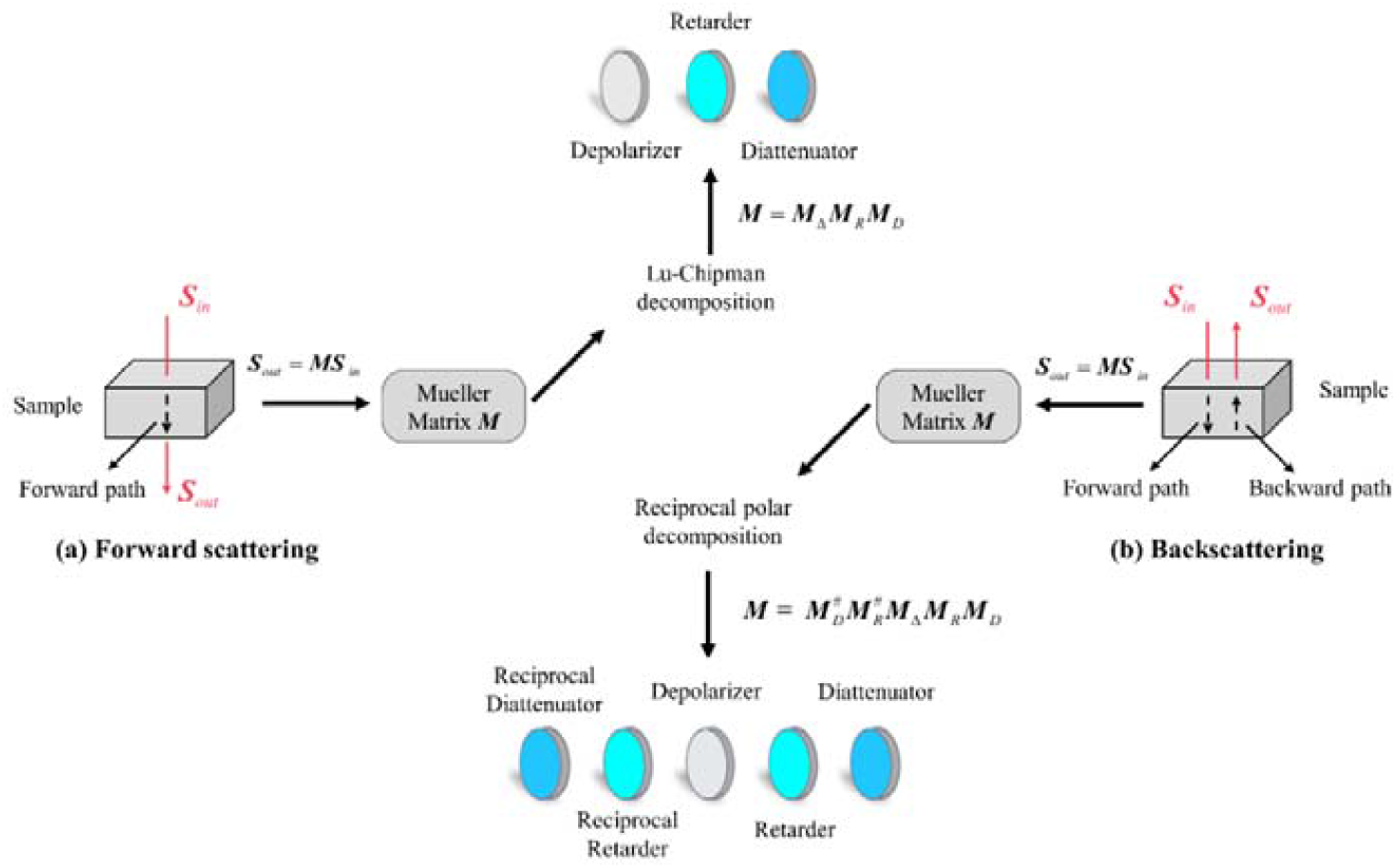
(a) Lu-Chipman decomposition of a forward scattering Mueller matrix. (b) Reciprocal polar decomposition of a backscattering Mueller matrix in reciprocal polarization imaging.

## 2 Reciprocal polarization imaging of complex media

Reciprocal polarization imaging relies on the reciprocity of optical systems in the absence of a magnetic field. We will denote the reciprocal matrix for a Mueller matrix ***M*** as ***M*** ^#^ for the optical system with the incident and outgoing beams interchanged. The reciprocal Mueller matrix can be written as ***M*** ^#^ = ***QM***^*T*^***Q*** where ***Q*** ≡ diag (1,1, −1,1) [19]. We introduce the reciprocal polar decomposition that decomposes a backscattering Mueller matrix into

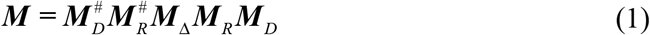

Where ***M***_*D*_, 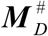 and ***M***_*R*_, 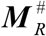 are, respectively, pairs of diattenuator matrices and retarder matrices in the forward and backward paths, and ***M***_Δ_ is the depolarizer matrix. The diattenuator and retarder matrices, ***M***_*D*_ and ***M***_*R*_, take the same form as in Lu-Chipman decomposition (see Supplementary information). Inside an exact backward geometry, the measured Mueller matrix ***M*** is the reciprocal of itself, i.e., ***M*** = ***QM***^*T*^ ***Q***. The depolarizer matrix further belongs to the type of the diagonal form [20,21] and reduces to ***M***_Δ_ = ***M***_Δ*d*_ ≡ diag (*d*_0_, *d*_1_, *d*_2_, *d*_3_). The reciprocal polar decomposition of the backscattering Mueller matrix ***M*** can be rewritten as:

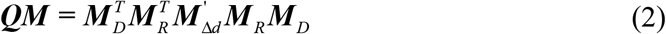

in terms of the symmetric matrix ***QM*** where 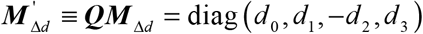. The diattenuator, retarder, and depolarizer matrices ***M***_*D*_, ***M***_*R*_, and ***M***_Δ_ of the sample are then determined by decomposition of Eq. (2) (see Methods).

Backscattering Mueller matrices are typically measured slightly off the normal direction to avoid specular reflection. In reciprocal polar decomposition of experimentally measured backscattering Mueller matrices, we replace ***QM*** by [***QM*** + (***QM***)^*T*^] / 2 to guarantee its symmetry. For comparison, results from Lu-Chipman decomposition are also shown. However, Lu-Chipman decomposition cannot be applied directly to the backscattering Mueller matrix as the incident and outgoing beams are defined according to different coordinate systems (see Fig. 1).

We replace the backscattering Mueller matrix ***M*** by ***M***_mirror_ ***M*** flipping the coordinate system for the outgoing beam to be the same as that of the incident beam before Lu-Chipman decomposition where ***M***_mirror_ = diag (1,1, −1, −1).

## 3 Results

### 3.1 Reciprocal polarization imaging of a birefringence resolution target

We first imaged NBS 1963A Birefringence Resolution Target (R2L2S1B, Thorlabs) in both backward and forward geometries with our custom polarization imaging system (see Fig. 2). The target contains a liquid crystal polymer pattern sandwiched between two N-BK7 glass substrates and has minimal diattenuation. The extracted linear retardance and orientation angle as well as depolarization for the target by the reciprocal polar decomposition in the backward geometry and by the Lu-Chipman decomposition in both the backward and forward geometries are compared (see Fig. 3). In addition, the horizontal profiles of the target orientation angle, linear retardance, and depolarization along the white line in Fig. 3 are shown in Fig. 4. The mean and standard deviation of the orientation angle, linear retardance, and depolarization for the birefringent (outlined by a red rectangle) and clear (outlined by a white rectangle) regions of the target measured in the forward and backward geometries are summarized in Table 1.

**Fig. 2.**
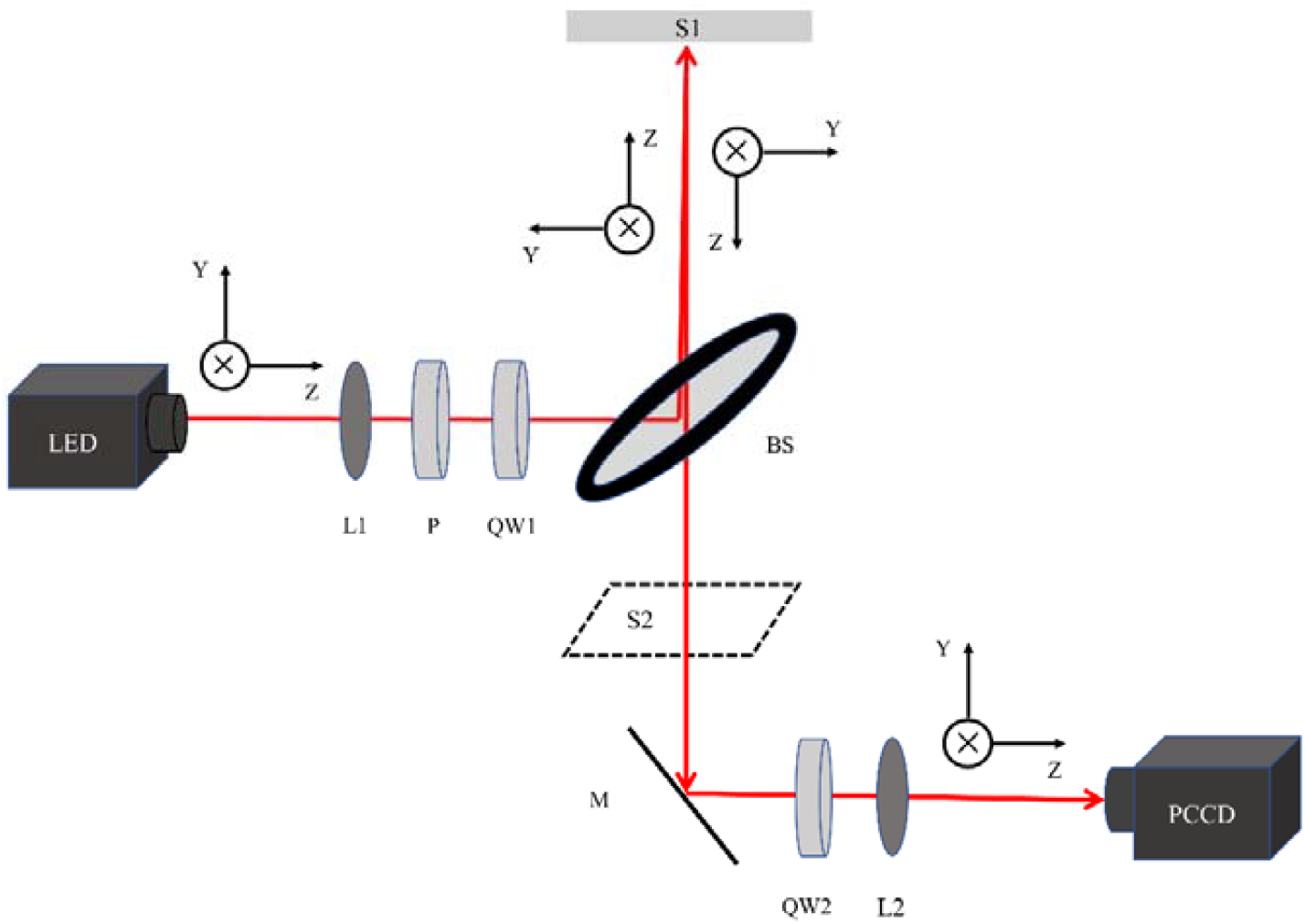
The polarization imaging system. P: polarizer; QW: quarter wave plates; L: lens; BS: beam splitter; M: mirror; PCCD: polarization camera; S1: sample position for backscattering Mueller matrix measurement; S2: sample position for forward-scattering Mueller matrix measurement.

**Fig. 3.**
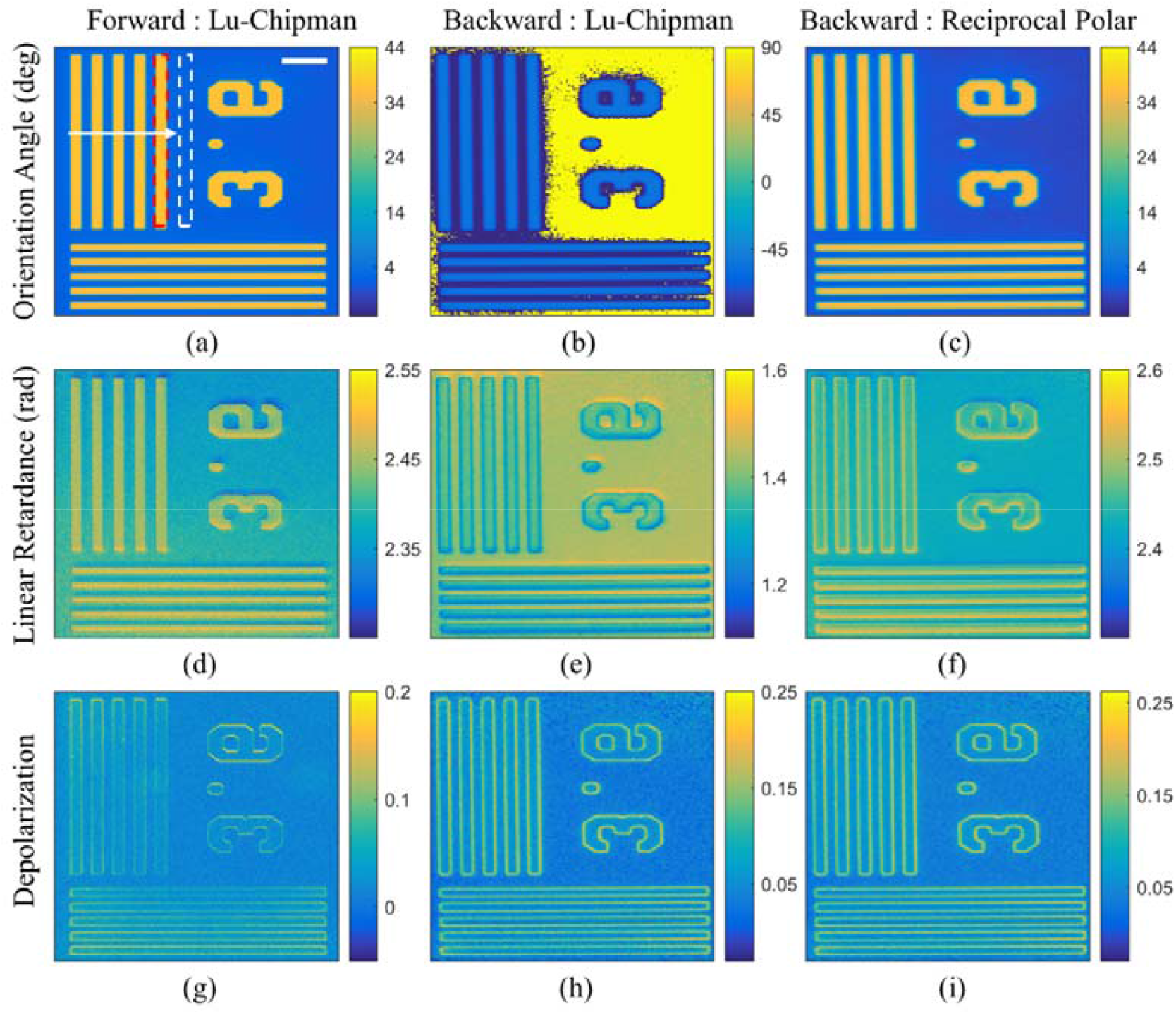
Polarization imaging of a birefringence resolution target in forward and backward geometries. The orientation angle, linear retardance, and depolarization from (a, d, g) Lu-Chipman decomposition of the Mueller matrix measured in the forward geometry; (b, e, h) Lu-Chipman decomposition of the Mueller matrix measured in the backward geometry; and (c, f, I) Reciprocal polar decomposition of the Mueller matrix measured in the backward geometry. Space bar: 0.5 mm.

**Fig. 4.**
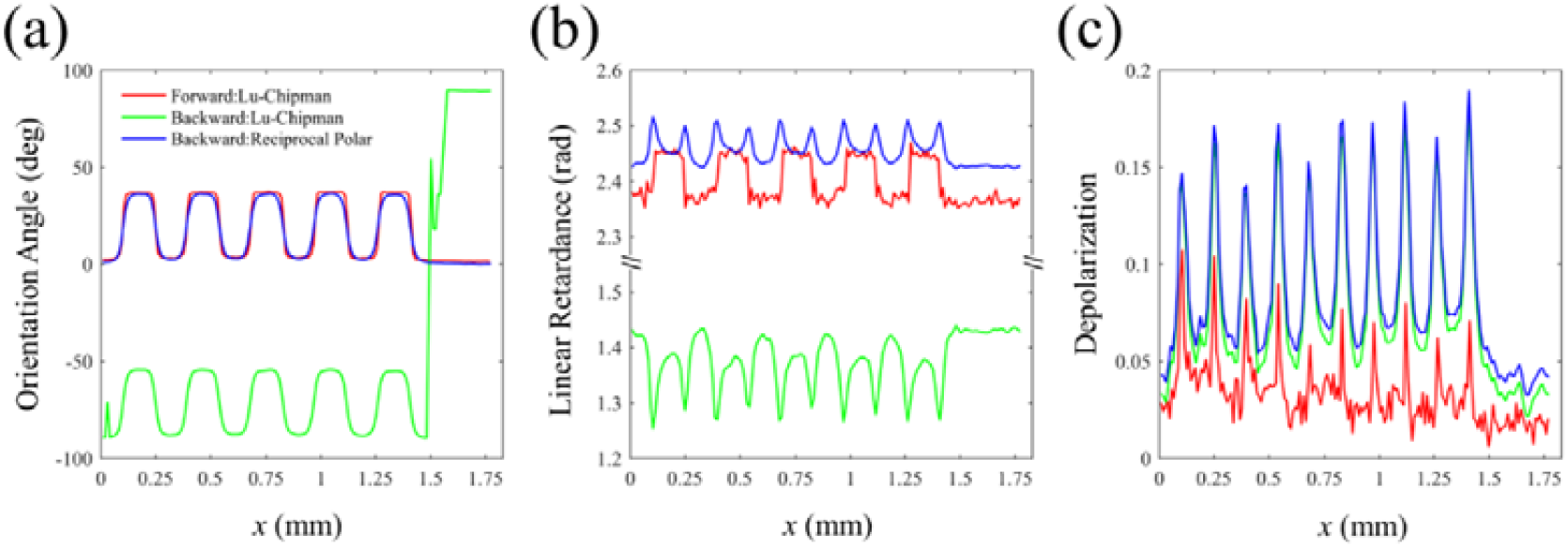
Profiles of (a) the target orientation angle, (b) linear retardance, and (c) depolarization from Lu-Chipman decomposition of the Mueller matrix measured in the forward geometry; Lu-Chipman decomposition of the Mueller matrix measured in the backward geometry; and reciprocal polar decomposition of the Mueller matrix measured in the backward geometry.

**Table 1.** Mean and standard deviation of the orientation angle, linear retardance, and depolarization for the birefringent (subscript ““B””) and clear (subscript ““C””) regions of the target measured in the forward and backward geometries.

| Parameters | Forward Geometry | Backward Geometry |  |
| --- | --- | --- | --- |
|  | Lu-Chipman | Lu-Chipman | Reciprocal Polar |
| $\theta_B$ (deg) | 36.67 (0.59) | -59.85 (7.70) | 35.35 (0.49) |
| $\theta_C$ (deg) | 1.67 (0.33) | 85.67 (25.46) | 0.10 (0.28) |
| $\delta_B$ (rad) | 2.448 (0.017) | 1.344 (0.038) | 2.471 (0.019) |
| $\delta_C$ (rad) | 2.370 (0.024) | 1.436 (0.010) | 2.429 (0.005) |
| $\Delta_B$ | 0.032 (0.018) | 0.093 (0.036) | 0.100 (0.036) |
| $\Delta_C$ | 0.016 (0.015) | 0.032 (0.010) | 0.041 (0.009) |

The recovered polarization parameters from the reciprocal polar decomposition of the backscattering Mueller matrix are in excellent agreement with those obtained from the Lu-Chipman decomposition of the Mueller matrix measured in the forward geometry other than a stronger depolarization in the former owing to the different detection geometry. The Lu-Chipman decomposition on the backscattering Mueller matrix fails to obtain the correct orientation angle and linear retardance. In particular, the orientation angle obtained by the Lu-Chipman decomposition of the backscattering Mueller matrix is off by 90 degrees and contains sporadic artifacts. The edges of the birefringent regions exhibit higher retardance and depolarization in the backscattering measurement (see Figs. 3 (f, i) and 4). This difference originates from edge diffraction, which leads to increased depolarization and larger retardance of light rays that transverse a longer path within the target. In contrast, the edges of the birefringent regions in the forward geometry only show slightly increased depolarization as the edge diffraction rays are much more dominant in the backscattering geometry where the specular reflection is removed than in the forward geometry. The obtained orientation and linear retardance by the Lu-Chipman decomposition of the forward scattering Mueller matrix and the reciprocal polar decomposition of the backscattering Mueller matrix agree with the data provided by the manufacturer (see Supplementary information).

### 3.2 Reciprocal polarization imaging of tissue

We then imaged a fresh beef section (thickness 100 μm) in both backward and forward geometries. The extracted tissue birefringence orientation angle, linear retardance, depolarization, and linear depolarization anisotropy (defined as *A ≡* (|*d*_1_| − |*d*_2_|)/(|*d*_1_|+s|*d*_2_|) by the reciprocal polar decomposition in the backward geometry and by the Lu-Chipman decomposition in both the backward and forward geometries are compared (see Fig. 5). The results measured in the backward geometry resemble those of the forward geometry allowing inevitable deformation of the tissue section between the two measurement geometries. The orientation angle from the reciprocal polar decomposition of the backscattering Mueller matrix is in closer agreement than that computed by the Lu-Chipman decomposition with the orientation angle measured in the forward geometry (mean squared error: 108 vs. 139, and correlation coefficient: 0.34 vs. 0.28). The red rectangle marks a region that Lu-Chipman decomposition of the backscattering Mueller matrix leads to significant errors in the orientation angle. The linear retardance from the Lu-Chipman decomposition of the backscattering Mueller matrix doubles that obtained by the reciprocal polar decomposition as it accounts for both forward and backward paths. Light depolarization is larger in the backscattering geometry than in the forward geometry. More importantly, tissue depolarization and depolarization anisotropy are distorted in the Lu-Chipman decomposition of the backscattering Mueller matrices. The images of tissue depolarization and depolarization anisotropy from the reciprocal polar decomposition of the backscattering Mueller matrix are much sharper than those obtained by the Lu-Chipman decomposition on the same data (sharpness: 2.43×10^4^ vs. 1.5×10^4^ for tissue depolarization and 1.71×10^6^ vs. 1.71×10^5^ for depolarization anisotropy).

**Fig. 5.**
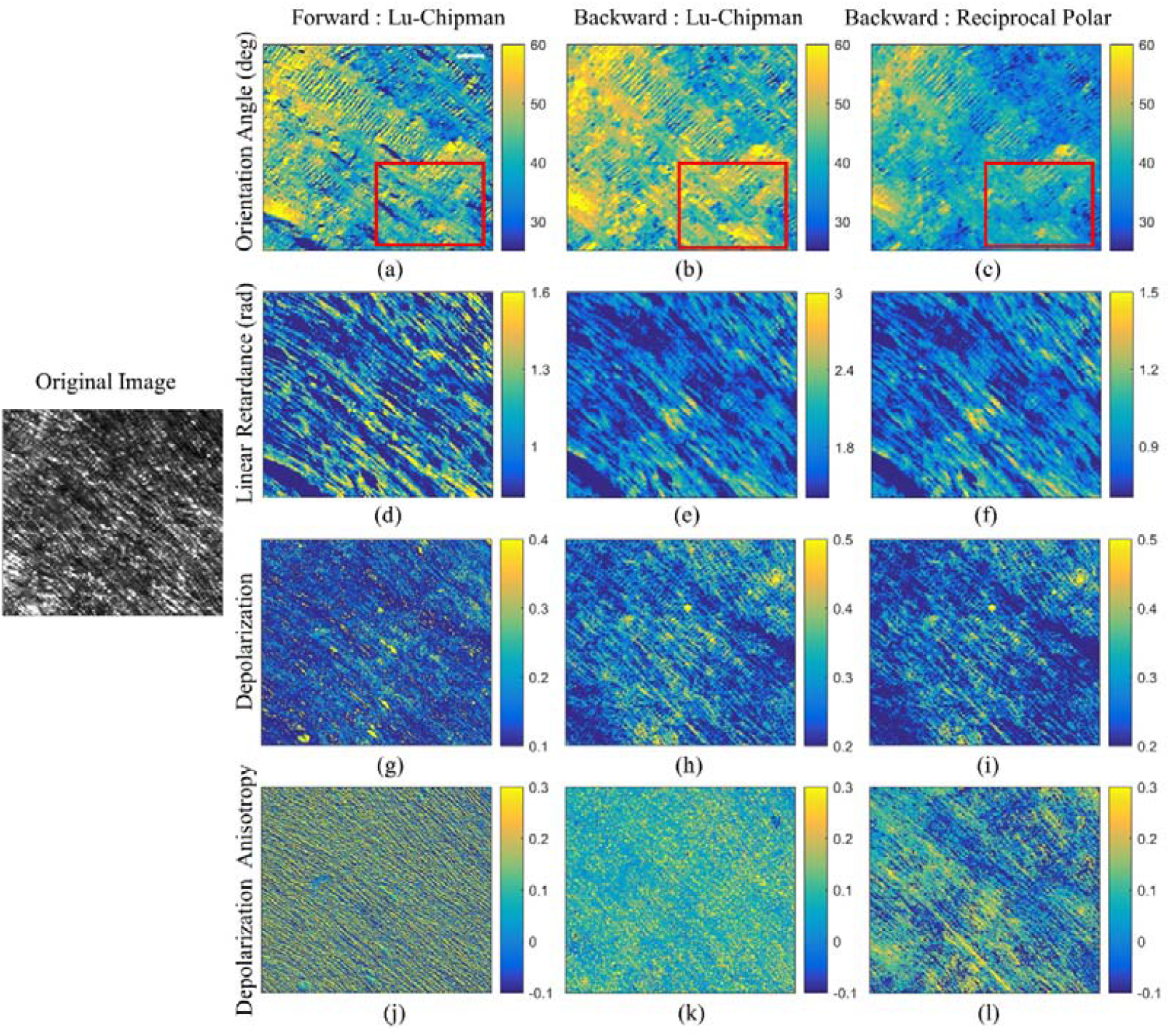
Polarization imaging of tissue in forward and backward geometries. The orientation angle, linear retardance, depolarization, and depolarization anisotropy from (a, d, g, j) Lu-Chipman decomposition of the Mueller matrix measured in the forward geometry; and (b, e, h, k) Lu-Chipman decomposition and (c, f, i, l) reciprocal polar decomposition of the Mueller matrix measured in the backward geometry. The left inset is the photo of the sample under unpolarized light illumination. Space bar: 0.5 mm.

Fig. 6 shows the corresponding images to Fig. 5 with fibers highlighted. The fiber region is selected for pixels of the top *p* percent depolarization values (*p*=65%). One representative fiber is outlined by a dashed white line with its normal direction (marked by white arrows) and the fiber orientation angle (marked by red arrows) computed by different decomposition methods. The mean and standard deviation of the orientation angle, linear retardance, depolarization, and depolarization anisotropy measured in the forward and backward geometries are summarized in Table 2 for all fibers as well as the one highlighted fiber. The highlighted fiber orientation angle (39.74 ± 0.61°) from reciprocal polar decomposition of the backscattering Mueller matrix is closer to the expected direction (41.31 ± 0.92°) normal to the fiber than that (44.40 ± 0.42°) obtained by the Lu-Chipman decomposition. Furthermore, the depolarization anisotropy recovered by the reciprocal polar decomposition of the backscattering Mueller matrix agrees well with that obtained by the Lu-Chipman decomposition of the Mueller matrix measured in the forward geometry. In contrast, the Lu-Chipman decomposition of the backscattering Mueller matrix yields erroneous depolarization anisotropy.

**Table 2.**
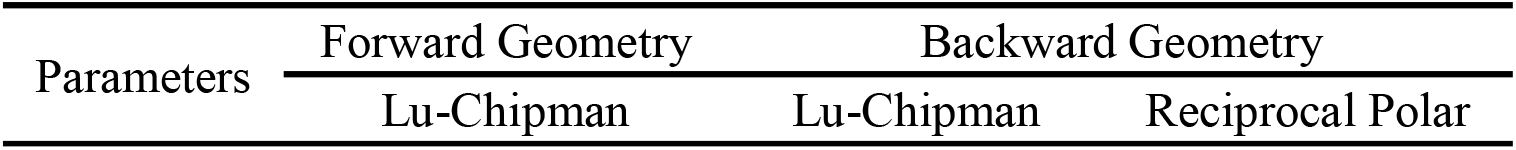
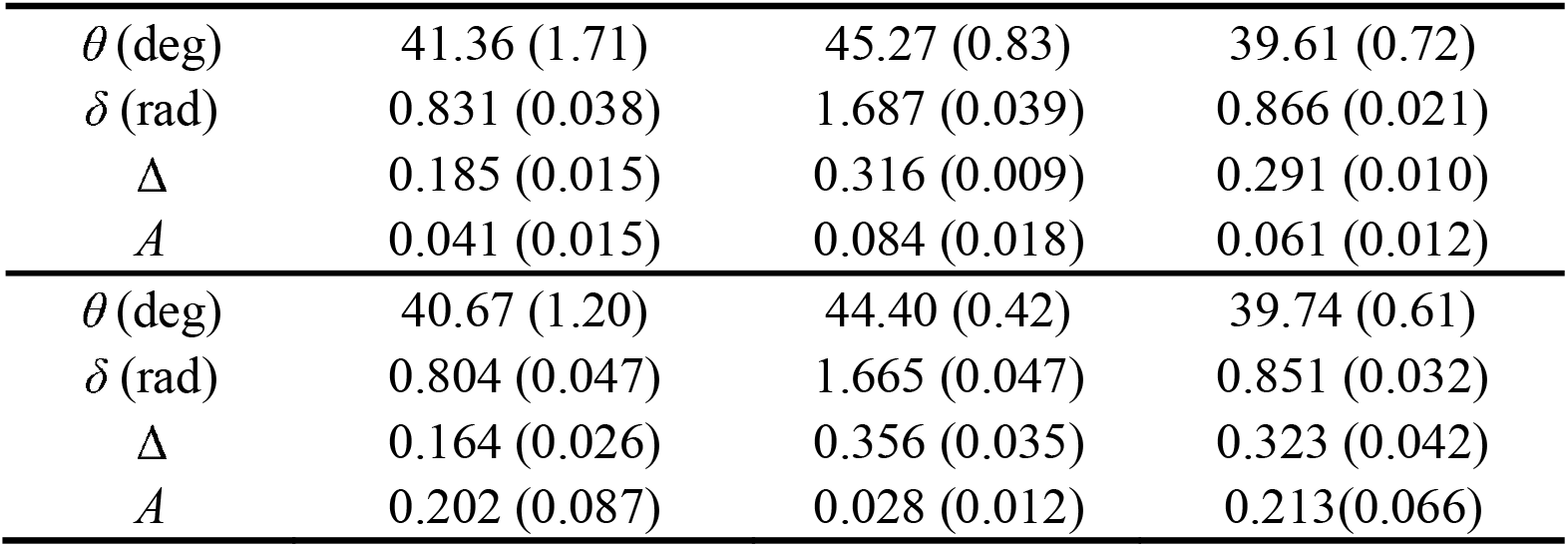
Mean and standard deviation of the fiber orientation angle, linear retardance, depolarization, and depolarization anisotropy measured in the forward and backward geometries for all fibers (top) and the one representative fiber (bottom).

| Parameters | Forward Geometry | Backward Geometry |  |
| --- | --- | --- | --- |
|  | Lu-Chipman | Lu-Chipman | Reciprocal Polar |
| $\theta$ (deg) | 41.36 (1.71) | 45.27 (0.83) | 39.61 (0.72) |
| $\delta$ (rad) | 0.831 (0.038) | 1.687 (0.039) | 0.866 (0.021) |
| $\Delta$ | 0.185 (0.015) | 0.316 (0.009) | 0.291 (0.010) |
| $A$ | 0.041 (0.015) | 0.084 (0.018) | 0.061 (0.012) |
| $\theta$ (deg) | 40.67 (1.20) | 44.40 (0.42) | 39.74 (0.61) |
| $\delta$ (rad) | 0.804 (0.047) | 1.665 (0.047) | 0.851 (0.032) |
| $\Delta$ | 0.164 (0.026) | 0.356 (0.035) | 0.323 (0.042) |
| $A$ | 0.202 (0.087) | 0.028 (0.012) | 0.213(0.066) |

**Fig. 6.**
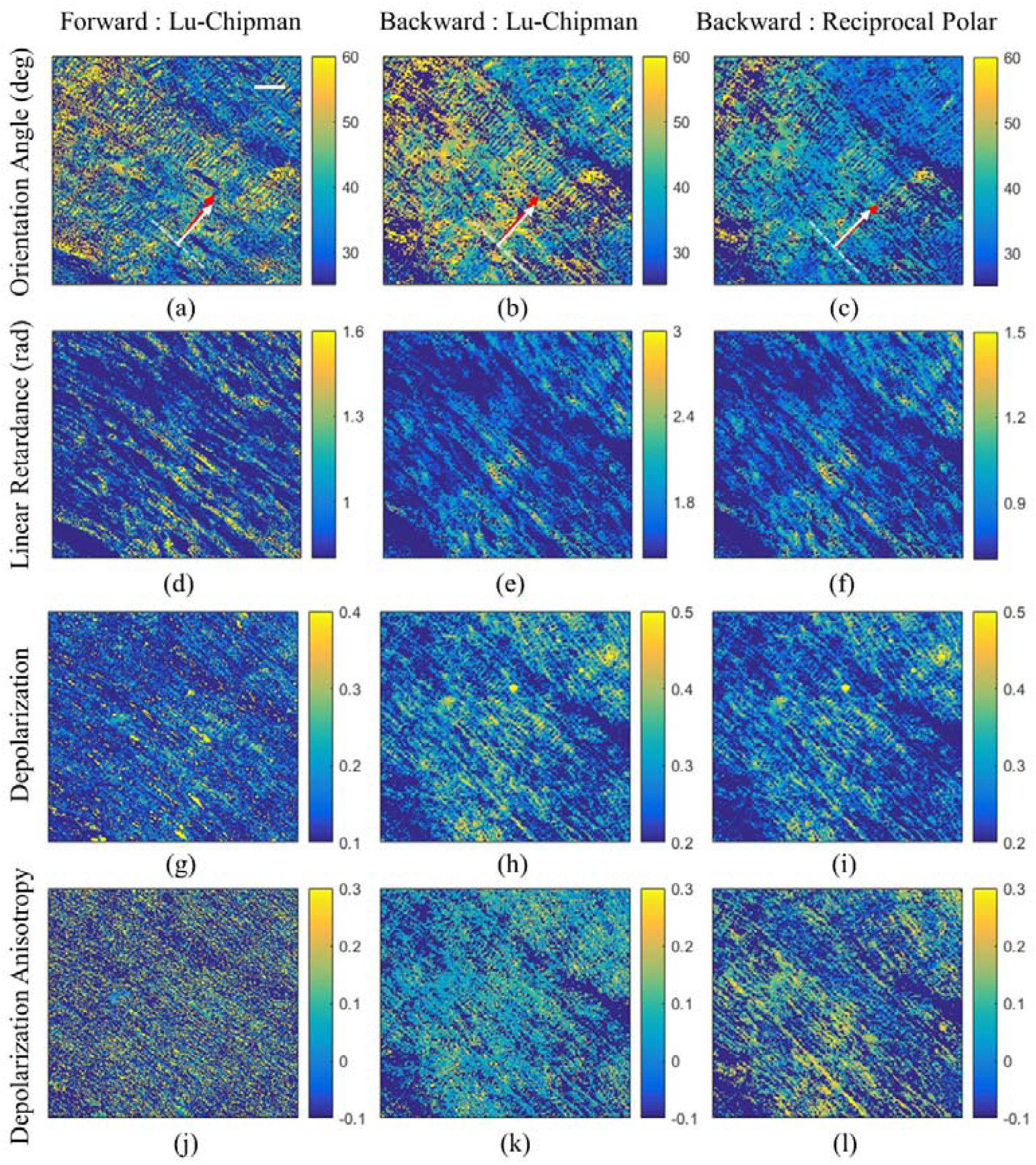
Polarization imaging of tissue in forward and backward geometries with fibers highlighted. The orientation angle, linear retardance, depolarization, and depolarization anisotropy from (a, d, g, j) Lu-Chipman decomposition of the Mueller matrix measured in the forward geometry; and (b, e, h, k) Lu-Chipman decomposition and (c, f, i, l) reciprocal polar decomposition of the Mueller matrix measured in the backward geometry. Space bar: 0.5 mm

### 3.3 Reciprocal polarization imaging of a chiral medium

Experimentally measured Mueller matrices for a chiral turbid medium (scattering coefficient μ_s_ =0.6 mm^-1^, anisotropy factor g=0.91, glucose concentration 5 M, and thickness 10 mm) in the forward and backward scattering geometries reported by Manhas[8]et al. was analyzed to demonstrate the applicability of reciprocal polarization imaging for chiral media. This medium is nonbirefringent, and the apparent linear retardance is attributed to light scattering[8]. The reciprocal polar decomposition shows the increase of linear retardance δ and optical rotation Ψ at a similar rate in the backward vs. forward scattering geometries, consistent with larger linear retardance and optical rotation in the backward scattering geometry owing to the increased path lengths for the detected photons. In contrast, the Lu-Chipman decomposition of the backscattering Mueller matrix yields an erroneous optical rotation in the backscattering geometry (see Table 3). Furthermore, the much longer path lengths of the detected photons further lead to increased light depolarization recovered from Lu-Chipman and reciprocal polar decompositions in the backward than forward geometries.

**Table 3.** Reciprocal polar decomposition of the Mueller matrix measured in the backward geometry for a chiral turbid medium[8] and the extracted polarization parameters compared to the Lu-Chipman decomposition of the Mueller matrices measured in the forward and backward geometries for the same medium.

| Backscattering $\mathbf{M}$ | | | | | | | | | | | | | | | |
| --- | --- | --- | --- | --- | --- | --- | --- | --- | --- | --- | --- | --- | --- | --- | --- |
| $\begin{pmatrix} 1 & -0.115 & -0.066 & 0.023 \\ -0.111 & 0.759 & -0.061 & -0.001 \\ -0.018 & 0.151 & -0.435 & -0.139 \\ -0.046 & 0.006 & 0.128 & -0.334 \end{pmatrix}$ | | | | | | | | | | | | | | | |
| | | | | $\mathbf{M}_\Delta$ | | | | $\mathbf{M}_R$ | | | | $\mathbf{M}_D$ | | | |
| | | | | $\begin{pmatrix} 0.997 & 0 & 0 & 0 \\ 0 & 0.790 & 0 & 0 \\ 0 & 0 & -0.425 & 0 \\ 0 & 0 & 0 & -0.359 \end{pmatrix}$ | | | | $\begin{pmatrix} 1 & 0 & 0 & 0 \\ 0 & 0.952 & -0.305 & -0.035 \\ 0 & 0.307 & 0.937 & 0.166 \\ 0 & -0.018 & -0.168 & 0.986 \end{pmatrix}$ | | | | $\begin{pmatrix} 1 & -0.066 & -0.020 & -0.013 \\ -0.066 & 1 & 0.001 & 0.000 \\ -0.020 & 0.001 & 0.998 & 0.000 \\ -0.013 & 0.000 & 0.000 & 0.998 \end{pmatrix}$ | | | |

| Parameters | Forward Geometry | Backward Geometry |  |
| --- | --- | --- | --- |
|  | Lu-Chipman | Lu-Chipman | Reciprocal Polar |
| d | 0.064 | 0.135 | 0.067 |
| $\Delta$ | 0.033 | 0.473 | 0.475 |
| $\delta$ (rad) | 0.055 | 0.336 | 0.170 |
| $\Psi$ (rad) | 0.069 | 0.034 | 0.157 |

## 4 Discussion and Conclusion

The Mueller matrix decomposition plays a central role in interpreting the polarized light-medium interaction and relating the evolution of the vector light wave to the physical property of a complex medium. Similar to polarization imaging that relies on the Lu-Chipman decomposition for factoring the forward-scattering Mueller matrix into a product of three matrices describing the medium depolarization, birefringence, and dichroism, reciprocal polarization imaging achieves the feat by the reciprocal polar decomposition of Mueller matrices measured in the backward scattering geometry. It factors the backscattering Mueller matrix into a product of **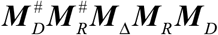** with the diattenuation and retardance along the backward path specified by the reciprocal of their counterparts 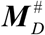 and 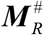 along the forward path based on the reciprocity of the optical wave in the forward and backward paths. Compared to the forward-scattering Mueller matrices, the backscattering Mueller matrices has reduced degrees of freedom of ten (***QM*** is a symmetric matrix). These ten degrees of freedom exactly map to ten polarization parameters: the diattenuation vector ***D***, the retardance vector (i.e., the linear retardance δ, its orientation θ, and the optical rotation Ψ), and the depolarization factors *d*_0_, *d*_1_, *d*_2,_ and *d*_3_. Reciprocal polarization imaging realizes a straightforward interpretation of the polarization measurement in the backward geometry and provides the determination of the same set of ten polarization parameters typically extracted from Mueller matrices measured in the forward geometry. It presents a significant advantage as the backward geometry is often preferred or the only feasible approach for bulk samples and more convenient in many applications.

We have demonstrated reciprocal polarization imaging with various complex non-chiral and chiral media and have shown the superiority of the reciprocal polar decomposition to the Lu-Chipman decomposition of backscattering Mueller matrices. The Lu-Chipman decomposition of the backscattering Mueller matrices produces similar yet distorted retardance, depolarization, and depolarization anisotropy images compared to the reciprocal polar decomposition for media of a low retardance (see Figs. 5 and 6). In particular, the Lu-Chipman decomposition fails to obtain the correct depolarization anisotropy from the backscattering Mueller matrices, whereas the reciprocal polar decomposition succeeds. For media of retardance exceeding π/2, significant errors in the orientation angle and retardance are further observed in the Lu-Chipman decomposition of the backscattering Mueller matrices (see Figs. 3 and 4). Additional examples of the failure of Lu-Chipman decomposition of backscattering Mueller matrices are given in Supplementary information. In all cases, the polarization properties of complex media determined by the reciprocal polar decomposition of the backscattering Mueller matrices are in excellent agreement with those obtained by polarization imaging of the same sample in the forward geometry. Furthermore, when light depolarization (Δ) is elevated, the accuracy of recovered media polarization parameters deteriorates from polarization imaging [22]. The performance of reciprocal polar decomposition for complex media of high depolarization warrants further study and will be published later.

Polarization imaging with Mueller matrix decomposition is a powerful means to quantify the diattenuation, retardance, and depolarization of complex media and link to the underlying microstructure and anisotropy. The measurement of the spatially resolved polarization properties has a wide array of applications in non-invasive characterization and diagnosis of complex random media[7,8], such as biological cells and tissue[9,10], for clinical and preclinical applications[4,12]. However, such measurement has traditionally been limited to transmission geometry. As backward geometry is ubiquitous and often preferred in applications from remote sensing to biomedical imaging, reciprocal polarization imaging will be instrumental in imaging complex media and opening up new applications of polarization optics in reflection geometry.

## 5 Materials and Methods

### 5.1 Reciprocal polar decomposition

The decomposition for Eq. (2) is performed similar to the symmetric decomposition [16] but much more straightforwardly. Noting

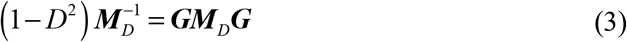

when the Mueller matrix ***M*** does not represent a perfect analyzer (*D* < 1) [14] where ***G*** ≡ diag (1, −1, −1, −1) is the Minkowski metric matrix, we have

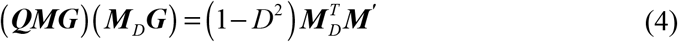

with

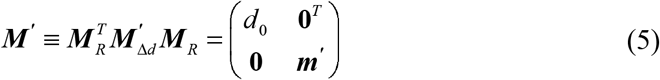

Expanding Eq. (4) after substituting Eqs. (S2) and (5) for respectively, lead to ***M***_*D*_ and ***M ^‘^***, respectively, lead to

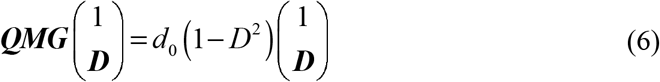

This equation has one unique positive eigenvalue (= *d*_0_ (1− *D*^2^)) and the associated eigenvector[16] providing the diattenuation vector ***D*** satisfying |***D***| = *D* < 1. With the determination of the diattenuation vector ***D***, the diattenuator matrix ***M***_***D***_ is obtained.

Afterward, Eq. (2) reduces to

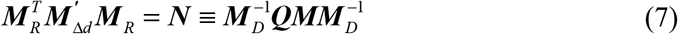

and equivalently

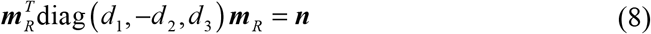

for the bottom right 3×3 submatrices ***m***_*R*_ and ***n*** of the matrices ***M***_*R*_ and ***N***. An orthogonal decomposition on ***n*** can then be used to determine ***m***_*R*_ and *d*_*i*_ (*i* = 1, 2,3). In addition, the order and the sign of the eigenvectors should be determined with *a priori* information, if available, regarding the depolarization properties of the medium. A convention of ordering the eigenvectors, which may be multiplied by ±1, to have a minimal total retardance is adopted in the absence of *a priori* information, similar to the Lu-Chipman and symmetric decompositions[14,23].The reciprocal polar decomposition of the backscattering Mueller matrix ***M*** into products (1) and (2) is then obtained.

We note that the degeneracy of the eigenvalues, *d*_1_, –*d*_2_, and *d*_3_, will render either part or complete loss of the determination of the retardance parameters as an arbitrary rotation can be applied to the degenerate eigenvectors. One potential degeneracy in the reciprocal polar decomposition arises for a depolarizer matrix ***M***_Δ*d*_ = diag (*d*_0_, *d*_1_, −*d*_1_, *d*_3_), which is fortunately rare and will result in an indeterminate circular retardance.

### 5.2 Polarization imaging system

The polarization imaging system is shown in Fig. 2. Collimated light (λ =633 nm) passes through a polarization state generator consisting of a rotating polarizer and a rotating quarter-wave plate and illuminates the sample after reflection by a beam splitter. The backscattered light by the sample is reflected by a mirror and passes through a second rotating quarter-wave plate before being recorded by a polarization camera (BFS-U3-51S5P-C, FLIR). The second rotating quarter-wave plate and the linear polarizers along 0°,45°, 90°, and 135° directions of the polarization camera form the polarization state analyzer. The Mueller matrix for the sample is measured in both the backward (sample at S1) and forward (sample at S2 and an additional mirror at S1) geometries after the removal of the stray light contributions and calibration of the Mueller matrix of the imaging system itself (see Supplementary information). In the backward geometry, the specular reflection from the sample surface is removed by slightly tilting the sample surface (∼ 5°). This Mueller imaging system has a field of view of 1.8×2.0 cm^2^. The acquisition of one complete Mueller matrix of the sample takes one minute.

### 5.3 Materials

Silverside beef rich in muscle fibers was selected for the experiment. One 1×1×1 cm^3^ of tissue was mounted in OCT embedding compound (Sakura) and kept at –80 °C for 10 hours. Tissue sections were cut using a cryostat microtome (HM525, Thermofisher) and mounted on a cover slip.

## Acknowledgments

We thank for Lili Ma’s assistance in preparing fresh beef section samples.

## Data availability

Data underlying the results presented in this paper are not publicly available at this time but may be obtained from the authors upon reasonable request.

## Supplemental document

See Supplementary information for supporting content.

