## Supplementary information for "Reciprocal polarization imaging of complex media"

### Reciprocal polarization imaging of complex media: supplemental document

Zhineng Xie, Guowu Huang, Weihao Lin, Xin Jin, Xiafei Qian, and Min Xu*

**This file includes:**

Figs. S1 to S7
Table S1 to S6

#### 1. Lu-Chipman polar decomposition

The Lu-Chipman decomposition of Mueller matrices is briefly outlined here for clarity and comparison with the reciprocal polar decomposition. Lu-Chipman decomposition decomposes the Mueller matrix into a product of three matrices: a depolarizer matrix $\boldsymbol{M}_{\Delta}$, a retarder matrix $\boldsymbol{M}_R$ , and a diattenuator matrix $\boldsymbol{M}_D$ [14,24], i.e.,

$$\boldsymbol{M} = \boldsymbol{M}_{\Delta}\boldsymbol{M}_R\boldsymbol{M}_D \tag{S1}$$

Different order for the above product has been introduced, and Eq. (S1) is widely adopted because the family in which the diattenuator matrix comes ahead of the depolarizer matrix always leads to a physically realizable Mueller matrix[25] . Here the diattenuator matrix is a symmetric matrix given by

$$\boldsymbol{M}_D = \begin{pmatrix} 1 & \boldsymbol{D}^T \\ \boldsymbol{D} & \boldsymbol{m}_D \end{pmatrix} \tag{S2}$$

in which $\boldsymbol{m}_D = \sqrt{1-D^2}\boldsymbol{I} + \left(1-\sqrt{1-D^2}\right)\hat{\boldsymbol{D}}\hat{\boldsymbol{D}}^T$ , $\boldsymbol{I}$ is a $3 \times 3$ identity matrix, $\hat{\boldsymbol{D}}$ is the unit vector for the diattenuation vector $\boldsymbol{D} = M_{00}^{-1}(M_{01}, M_{02}, M_{03})^T$ of length $D$, and $M_{ij}$ $(i, j = 0,1,2,3)$ is the element at the $i$-th row and $j$-th column of the Mueller matrix $\boldsymbol{M}$. The length is assumed $D<1$ to ensure a non-singular matrix $\boldsymbol{M}_D$ (for $\boldsymbol{M}_D^{-1}$ to exist). The length $D=1$ corresponds to a perfect analyzer that $\boldsymbol{M}$ has an indeterminate decomposition[14] .

The retarder matrix is written as

$$\boldsymbol{M}_R = \begin{pmatrix} 1 & \boldsymbol{0}^T \\ \boldsymbol{0} & \boldsymbol{m}_R \end{pmatrix} \tag{S3}$$

where $\boldsymbol{m}_R$ is a positive definite three-dimensional rotation matrix. The retarder matrix $\boldsymbol{M}_R$ can further be expressed as an ordered product of a circular retarder of retardance $\Psi$ and a linear retarder of retardance $\delta$ with its fast axis oriented at $\theta$. Their values are given by

$$\delta = \cos^{-1}\left(\sqrt{\left((M_R)_{1,1} + (M_R)_{2,2}\right)^2 + \left((M_R)_{1,2} - (M_R)_{2,1}\right)^2} - 1\right) \tag{S4}$$

$$\theta = -\frac{1}{2}\operatorname{atan2}\left((M_R)_{3,1}, (M_R)_{3,2}\right) \tag{S5}$$

and

$$\Psi = \frac{1}{2}\operatorname{atan2}\left((M_R)_{1,2} - (M_R)_{2,1}, (M_R)_{1,1} + (M_R)_{2,2}\right) \tag{S6}$$

Furthermore, the total retardance and the depolarization coefficient are given by

$$R=\cos^{-1}\left[2\cos^2\Psi\cos^2\left(\frac{\delta}{2}\right)-1\right] \tag{S7}$$

and

$$\Delta=1-\frac{1}{3}\left[\mathrm{tr}\left|\frac{\boldsymbol{M}_\Delta}{(\boldsymbol{M}_\Delta)_{00}}\right|-1\right] \tag{S8}$$

#### 2. Calibration of the polarization imaging system

The polarization state generator (consisting of a rotating polarizer and a rotating quarter wave plate) generates four input polarization states: $0^\circ$ (Stokes vector $[1\quad 1\quad 0\quad 0]^T$), $45^\circ$ (Stokes vector $[1\quad 0\quad 1\quad 0]^T$), $90^\circ$ (Stokes vector $[1\quad -1\quad 0\quad 0]^T$) linear polarization, and circular polarization (Stokes vector $[1\quad 0\quad 0\quad 1]^T$) states. The polarization state analyzer of a rotating quarter wave plate and a polarization camera[26] uses the approach presented.

The calibration of the Mueller imaging system follows the steps outlined in Fig. S1. First, Stokes vector $\boldsymbol{S}_{0^\circ}$, $\boldsymbol{S}_{45^\circ}$, $\boldsymbol{S}_{90^\circ}$, and $\boldsymbol{S}_{circular}$ for the four input polarization states were measured using the configuration (a). Second, Mueller matrices of mirrors M1 and M2, $\boldsymbol{M}_{M1}$ and $\boldsymbol{M}_{M2}$, were measured using the configuration (b). Third, the reflection and transmission Mueller matrices of the beam splitter BS, $\boldsymbol{R}$ and $\boldsymbol{T}$, were measured in the configuration (c) and (d), respectively. The obtained values are given below:

$$\begin{aligned}
\boldsymbol{S}_{0^\circ}&=\begin{bmatrix}1 & 0.983 & 0.002 & -0.025\end{bmatrix}^T\\
\boldsymbol{S}_{45^\circ}&=\begin{bmatrix}0.984 & 0.019 & 0.965 & 0.020\end{bmatrix}^T\\
\boldsymbol{S}_{90^\circ}&=\begin{bmatrix}0.978 & -0.952 & 0.003 & -0.021\end{bmatrix}^T\\
\boldsymbol{S}_{circular}&=\begin{bmatrix}0.972 & 0.03 & 0.061 & 0.947\end{bmatrix}^T
\end{aligned} \tag{S9}$$

$$\boldsymbol{S}_{in}=\begin{bmatrix}\boldsymbol{S}_{0^\circ} & \boldsymbol{S}_{90^\circ} & \boldsymbol{S}_{45^\circ} & \boldsymbol{S}_{circular}\end{bmatrix} \tag{S10}$$

$$\boldsymbol{M}_{M1}=\begin{bmatrix}1 & 0.141 & 0.013 & -0.005\\ 0.135 & 0.986 & 0.001 & 0.033\\ -0.020 & 0.059 & -0.997 & -0.028\\ -0.032 & -0.028 & 0.068 & -0.999\end{bmatrix} \tag{S11}$$

$$\boldsymbol{M}_{M2}=\begin{bmatrix}1 & -0.053 & -0.027 & -0.009\\ -0.052 & 0.9956 & -0.025 & -0.021\\ 0.038 & -0.002 & -0.942 & 0.371\\ -0.002 & -0.073 & -0.380 & -0.884\end{bmatrix} \tag{S12}$$

$$\boldsymbol{R}=\begin{bmatrix}1 & 0.366 & -0.011 & -0.024\\ 0.344 & 0.998 & -0.078 & -0.038\\ -0.031 & -0.013 & -0.851 & -0.316\\ -0.033 & -0.034 & 0.281 & -0.814\end{bmatrix} \tag{S13}$$

$$
\boldsymbol{T} = \begin{bmatrix} 1 & -0.395 & 0.015 & -0.001 \\ -0.378 & 0.997 & 0.006 & -0.023 \\ 0.025 & -0.050 & 0.907 & 0.103 \\ 0.032 & -0.001 & -0.045 & 0.874 \end{bmatrix} \tag{S14}
$$

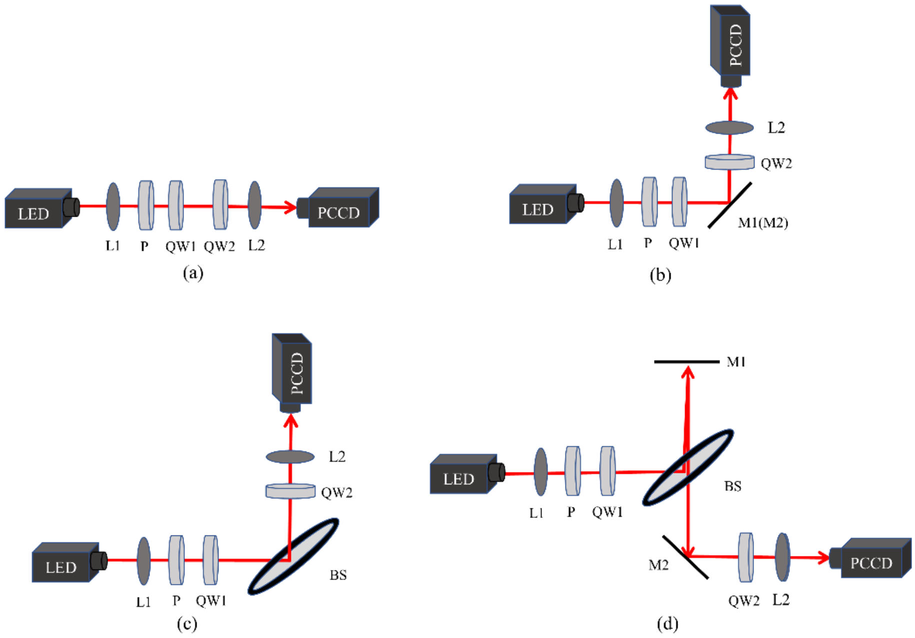


Fig. S1. Calibration of the polarization imaging system.

The measured Mueller matrix $\boldsymbol{M}_{out}$ is given by

$$
\boldsymbol{M}_{out} = \boldsymbol{M}_{M2}\boldsymbol{T}\boldsymbol{M}_{Sample}\boldsymbol{R}\boldsymbol{S}_{in} \tag{S15}
$$

in the backscattering geometry and is given by

$$
\boldsymbol{M}_{out} = \boldsymbol{M}_{M2}\boldsymbol{M}_{Sample}\boldsymbol{T}\boldsymbol{M}_{M1}\boldsymbol{R}\boldsymbol{S}_{in} \tag{S16}
$$

in the forward geometry, which contains an additional mirror M1 placed at the S1 position. The Mueller matrix $\boldsymbol{M}_{\text{sample}}$ for the sample is then solved from $\boldsymbol{M}_{out}$.

#### 3. Birefringent resolution target

The NBS 1963A Birefringence Resolution Target (Thorlabs) has 26 groups of lines, with five horizontal lines and five vertical lines in each group. The line group of 3.6 cycles/mm with a minimum period of 0.278 mm is selected for imaging in the experiment (see Fig. S2).

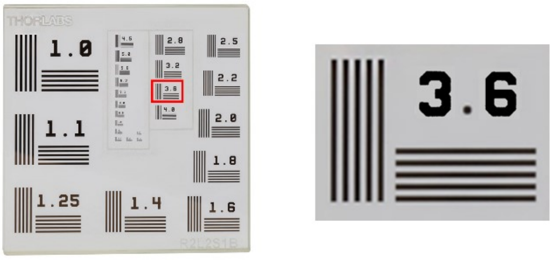


(a) (b)

Fig. S2. NBS 1963A Birefringence Resolution Target.

The polarization parameters of the selected horizontal line group provided by the manufacturer are summarized in Table S1 when the target is oriented as in the main text.

**Table S1.The polarization parameters provided by the manufacturer.**

| $\theta_B$ (deg) | $\theta_C$ (deg) | $\delta_B$ (rad) | $\delta_C$ (rad) |
|---|---|---|---|
| 34.5 ± 4.3 | 0.1 ± 3.8 | 2.436 ± 0.084 | 2.383 ± 0.068 |

#### 4. Measured orientation angle, linear retardance, and depolarization of the birefringent resolution target when placed in different directions

Measurements were performed for the target when placed in three different directions (#1: the horizontal line along the *x*-axis, #2: the target rotated 14.3° counterclockwise from #1, #3: the target rotated 13.3° clockwise from #1). The orientation angle, linear retardance, and depolarization of the target for #1, #2, and #3 from the Lu-Chipman decomposition of the forward scattering Mueller matrix, the Lu-Chipman decomposition, and the reciprocal polar decomposition of the backscattering Mueller matrix were obtained.

The orientation angle ($\theta$), linear retardance ($\delta$), and depolarization ($\Delta$) are shown in Figs. S3-S5. Their mean and standard deviation are shown in Table S2-S4. The orientation angles and the linear retardance obtained from the Lu-Chipman decomposition of the forward-scattering Mueller matrix and the reciprocal polar decomposition of the backscattering Mueller matrix are in excellent agreement. The orientation angles of the target for #1, #2, and #3 obtained by both the Lu-Chipman decomposition in the forward geometry and the reciprocal polar decomposition in the backward direction correctly follow their relative rotations. As expected, the depolarization of the backscattering Mueller matrix is larger than that of the forward scattering Mueller matrix.

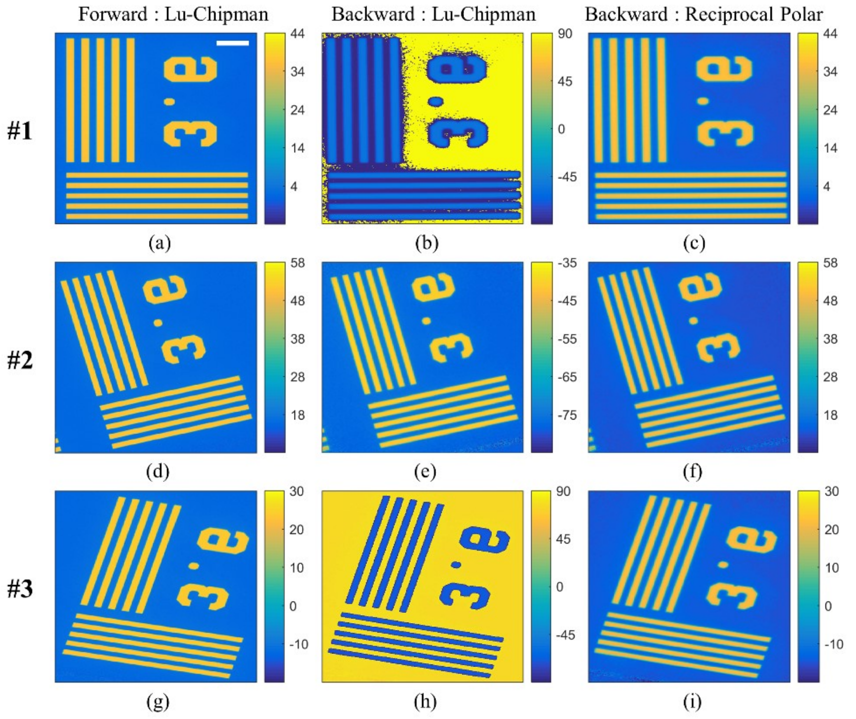


Fig. S3. Orientation angle (in degrees) of the target placed in different directions. Space bar: 0.5 mm.

Table S2. Mean and standard deviation of the orientation angle (θ). $\theta_B$ : Orientation angle of the birefringent region. $\theta_C$: Orientation angle of the clear region.

| | | Forward Geometry | Backward Geometry | |
|---|---|---|---|---|
| | | Lu-Chipman | Lu-Chipman | Reciprocal Polar |
| **#1** | $\theta_B$ (deg) | 36.67 (0.59) | -59.85 (7.70) | 35.35 (0.49) |
| | $\theta_C$ (deg) | 1.67 (0.33) | 85.67 (25.46) | 0.10 (0.28) |
| **#2** | $\theta_B$ (deg) | 50.81 (1.53) | -41.05 (0.92) | 49.07 (1.10) |
| | $\theta_C$ (deg) | 16.27 (0.33) | -76.84 (0.33) | 14.12 (0.29) |
| **#3** | $\theta_B$ (deg) | 23.47 (1.64) | -69.96 (0.82) | 20.83 (0.90) |
| | $\theta_C$ (deg) | -11.56 (0.32) | 76.05 (0.31) | -13.78 (0.27) |

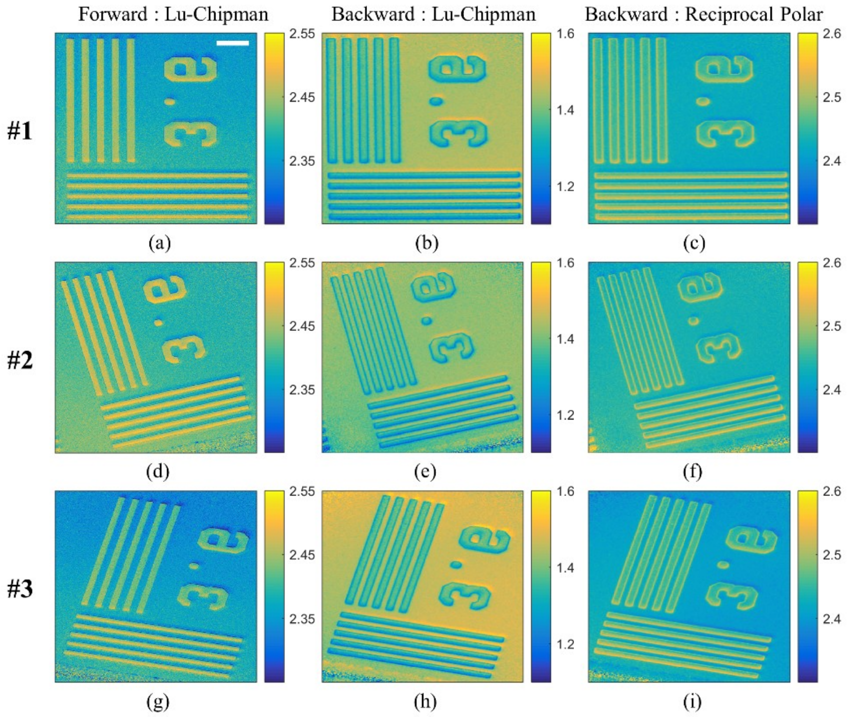


Fig. S4. Linear retardance (rad) of the target placed in different directions. Space bar: 0.5 mm.

Table S3. Mean and standard deviation of linear retardance (rad). $\delta_B$: Linear retardance of the birefringent region. $\delta_C$: Linear retardance of the clear region.

| | | Forward Geometry | Backward Geometry | |
|---|---|---|---|---|
| | | Lu-Chipman | Lu-Chipman | Reciprocal Polar |
| **#1** | $\delta_B$ (rad) | 2.448 (0.017) | 1.344 (0.038) | 2.471 (0.019) |
| | $\delta_C$ (rad) | 2.370 (0.024) | 1.436 (0.010) | 2.429 (0.005) |
| **#2** | $\delta_B$ (rad) | 2.465 (0.019) | 1.353 (0.045) | 2.467 (0.022) |
| | $\delta_C$ (rad) | 2.396 (0.021) | 1.417 (0.011) | 2.433 (0.005) |
| **#3** | $\delta_B$ (rad) | 2.419 (0.022) | 1.354 (0.034) | 2.466 (0.017) |
| | $\delta_C$ (rad) | 2.348 (0.024) | 1.467 (0.011) | 2.407 (0.005) |

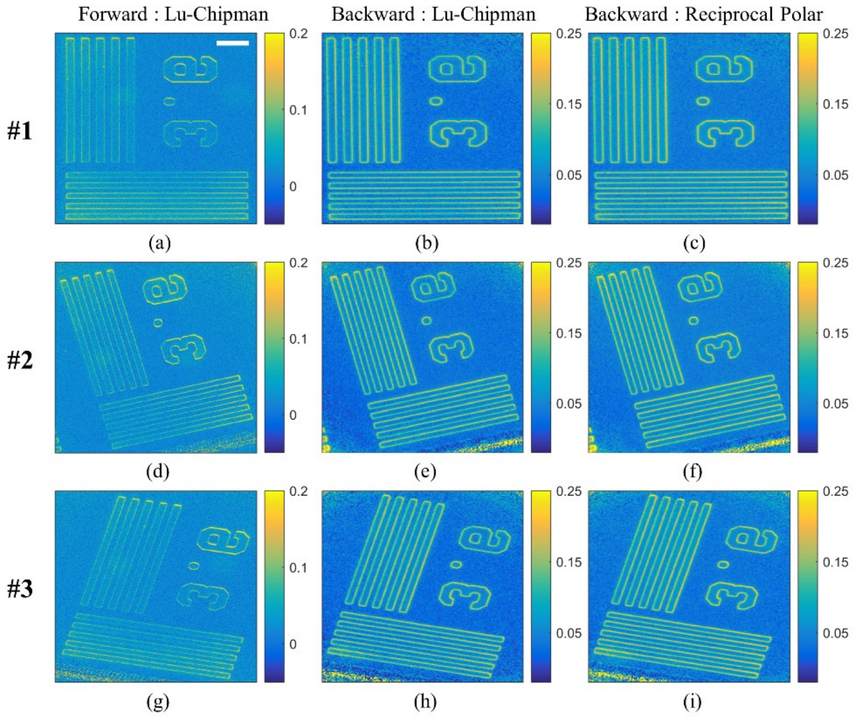


Fig.S5. Depolarization of the target placed in different directions. Space bar: 0.5 mm.

Table S4. Mean and standard deviation of depolarization of the target. $\Delta_B$: Depolarization of the birefringent region. $\Delta_C$: Depolarization of the clear region.

| | | Forward Geometry | Backward Geometry | |
|---|---|---|---|---|
| | | Lu-Chipman | Lu-Chipman | Reciprocal Polar |
| **#1** | $\Delta_B$ | 0.032 (0.018) | 0.093 (0.036) | 0.100 (0.036) |
| | $\Delta_C$ | 0.016 (0.015) | 0.032 (0.010) | 0.041 (0.009) |
| **#2** | $\Delta_B$ | 0.037 (0.024) | 0.095 (0.040) | 0.108 (0.041) |
| | $\Delta_C$ | 0.019 (0.016) | 0.034 (0.009) | 0.043 (0.009) |
| **#3** | $\Delta_B$ | 0.030 (0.018) | 0.092 (0.041) | 0.098 (0.039) |
| | $\Delta_C$ | 0.017 (0.016) | 0.028 (0.011) | 0.036 (0.010) |

#### 5. Failure of Lu-Chipman decomposition of backscattering Mueller matrices

We provide simulated examples showing the failure of the Lu-Chipman decomposition of backscattering Mueller matrices. The first example is a backscattering Mueller matrix with a diattenuation vector $\boldsymbol{D}=\left(0.1,0.1,0.1\right)^T$, linear retardance $\delta$=60°, its orientation $\theta$=−89°, optical rotation $\Psi$=6°, and depolarization factors $d_0$=1, $d_1$=1, $d_2$=−0.8, and $d_3$=−0.6. Figure S6 and Table S5 show the results of Reciprocal polar decomposition and Lu-Chipman decomposition, respectively. The second example is identical to the first one, except the linear retardance $\delta$ is modified from 60° to 100°. Figure S7 and Table S6 show the corresponding results.

In both examples, the parameters recovered by reciprocal polar decomposition are in excellent agreement with the ground truth. In contrast, the Lu-Chipman decomposition yields erroneous values for the diattenuation vector, the linear retardance, the optical rotation, $d_2$, $d_3$, and depolarization anisotropy A. It also produces erroneous orientation $\theta$ for the retardance exceeding π/2. The behavior of the simulated data is consistent with the results reported in the main text for the birefringence resolution target, the tissue sample, and the chiral medium.

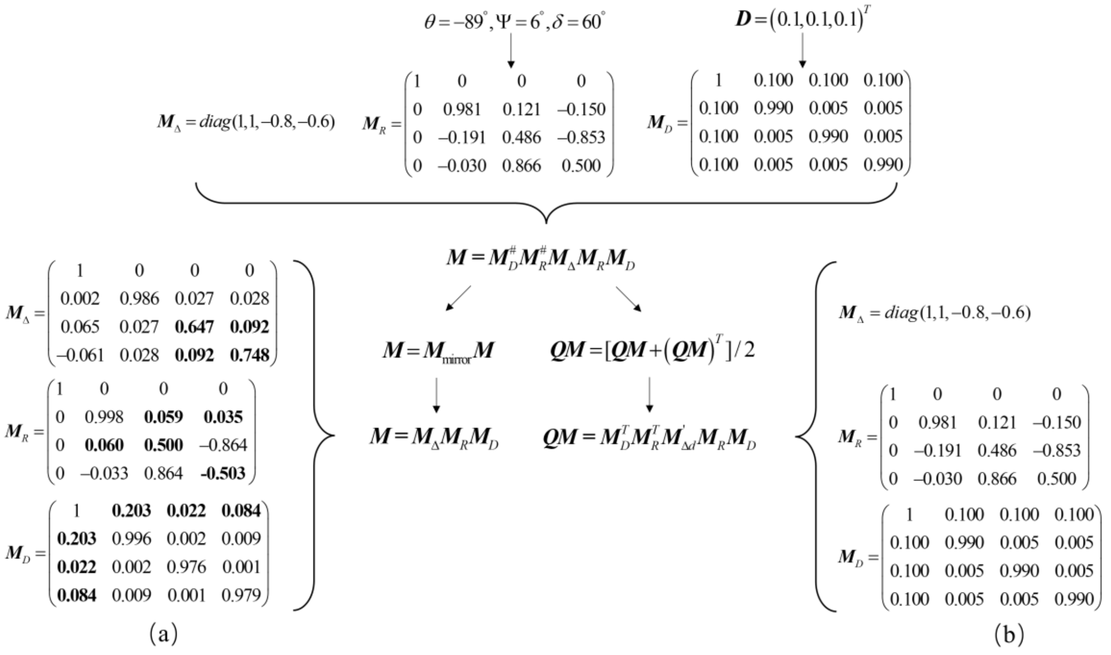

Fig. S6. Decomposition of simulated backscattering Mueller matrices. (a) Lu-Chipman decomposition, and (b) Reciprocal polar decomposition. Erroneous values are highlighted in bold.

**Table S5. The polarization parameters of decomposition of backscattering Mueller matrices. Erroneous values are highlighted in bold.**

| Parameters | Backward Geometry | | Ground Truth |
|---|---|---|---|
| | Lu-Chipman | Reciprocal Polar | |
| $\boldsymbol{D}$ | $\mathbf{(0.20,0.02,0.08)^T}$ | $(0.10,0.10,0.10)^T$ | $(0.10,0.10,0.10)^T$ |
| $\theta$ (deg) | -88.9 | -89.0 | -89.0 |
| $\delta$ (deg) | **120.2** | 60.0 | 60.0 |
| $\Psi$ (deg) | **-0.1** | 6.0 | 6.0 |
| $\Delta$ | 0.21 | 0.20 | 0.20 |
| d1 | 0.99 | 1.00 | 1.00 |
| d2 | **0.65** | -0.80 | -0.80 |
| d3 | **0.75** | -0.60 | -0.60 |
| A | **0.21** | 0.11 | 0.11 |

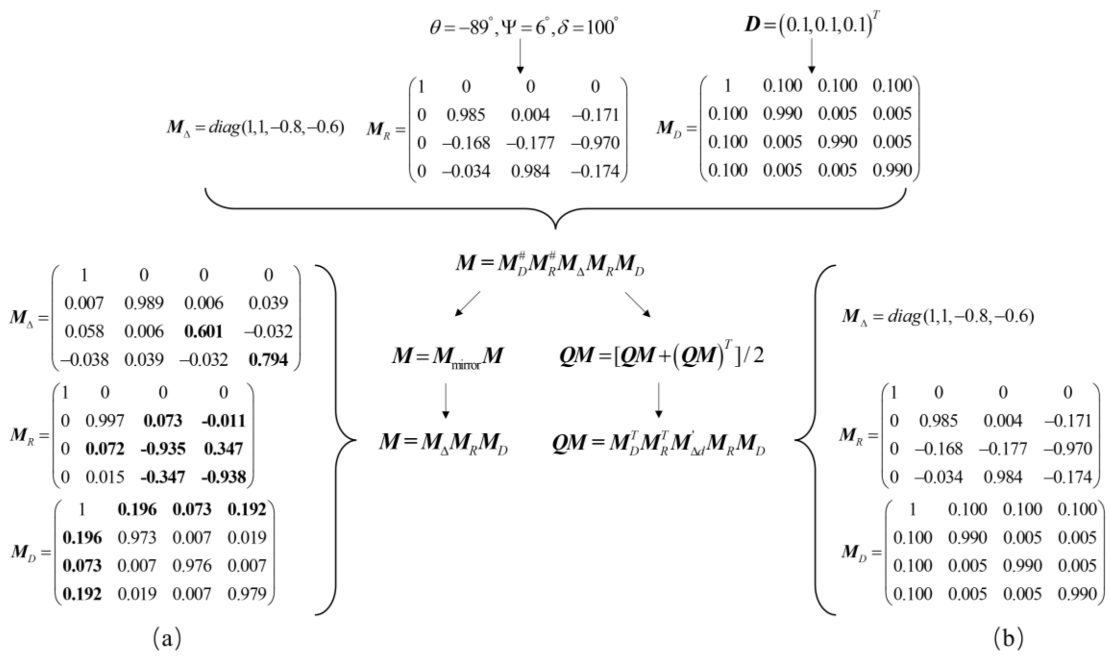


Fig. S7. Decomposition of simulated backscattering Mueller matrices with an altered linear retardance. (a) Lu-Chipman decomposition, and (b) Reciprocal polar decomposition. Erroneous values are highlighted in bold.

**Table S6. The polarization parameters of decomposition of backscattering Mueller matrices of an altered linear retardance. Erroneous values are highlighted in bold.**

| Parameters | Backward Geometry | | Ground Truth |
|---|---|---|---|
| | Lu-Chipman | Reciprocal Polar | |
| $\boldsymbol{D}$ | **(0.20,0.07,0.19)$^T$** | (0.10,0.10,0.10)$^T$ | (0.10,0.10,0.10)$^T$ |
| $\theta$ (deg) | **1.2** | -89.0 | -89.0 |
| $\delta$ (deg) | **159.7** | 100.0 | 100.0 |
| Ψ (deg) | **0.4** | 6.0 | 6.0 |
| Δ | 0.21 | 0.20 | 0.20 |
| d1 | 0.99 | 1.00 | 1.00 |
| d2 | **0.60** | -0.80 | -0.80 |
| d3 | **0.79** | -0.60 | -0.60 |
| A | **0.24** | 0.11 | 0.11 |
